# TDP-43 promotes efficient HSV-1 replication in human DRG-derived neurons

**DOI:** 10.1101/2025.07.08.662712

**Authors:** Shirley E. Braspenning, Denise Ohnezeit, Olivia A. DeGulis, Angus C. Wilson, Ian J. Mohr

## Abstract

TAR DNA-binding protein 43 (TDP-43) is a versatile nuclear RNA-binding protein that performs important functions in RNA localization, processing and stability. In the neurodegenerative disease amyotrophic lateral sclerosis (ALS) TDP-43 forms toxic, insoluble cytoplasmic aggregates that ultimately lead to neuronal loss. Although TDP-43 is expressed in every cell type, its function and subcellular localization are particularly important for neuronal homeostasis. However, it is unknown if TDP-43 has a role during herpesvirus infection. Herpes simplex virus type-1 (HSV-1), a ubiquitous neurotropic pathogen, is considered a contributing factor to neurodegenerative disorders. In this study, we tested the requirement for TDP-43 during HSV-1 infection in neuronal and non-neuronal cells. HSV-1 infection of epithelial cells and primary fibroblasts did not change overall TDP-43 abundance, nor did TDP-43 depletion detectably alter HSV-1 replication in a multicycle growth experiment. By contrast, when TDP-43 was depleted in neuronally derived, matured HD10.6 cells, HSV-1 infectious virus production was significantly reduced in both single- and multicycle growth experiments. Notably, TDP-43 depletion restricts viral lytic gene expression at the *immediate-early* phase. Through nanopore direct RNA-sequencing we uncovered enhanced intron retention in two essential viral genes upon TDP-43 depletion. Thus, while depletion of TDP-43 does not affect replication in epithelial cells and fibroblasts, TDP-43 is required for efficient replication in HD10.6 cells through modifying the abundance and splicing of viral mRNAs.

**IMPORTANCE:** Herpes simplex virus type-1 is a widespread neurotropic pathogen that can cause life-threatening infections of the brain and is increasingly linked to neurodegenerative disease. However, due to the lack of scalable *in vitro* human neuronal models or small animal models that recapitulate disease, little is known about virus-host interactions in neurons specifically. Using human epithelial cells, primary fibroblasts and a human neuron-derived cell line, we uncovered a cell type specific TDP-43 requirement for efficient HSV-1 virus replication. TDP-43 is a critical neuronal disease gene, and we showed it promotes virus gene expression and splicing of viral mRNAs in neuron-derived cells. This work provides valuable insights into the possible etiology of neurodegenerative disease and highlights the importance of studying virus-host interactions in relevant model systems.

## INTRODUCTION

TAR DNA-binding protein 43 (TDP-43) is a ubiquitously expressed nuclear RNA binding protein with key roles in RNA stability, processing and localization (1–3). Originally, TDP-43 was identified through its ability to bind the TAR DNA sequence of human immunodeficiency virus type 1 (HIV-1) (4), and restrict HIV-1 transcription. Subsequent research identified roles for TDP-43 in regulation of transcription (5, 6), miRNA biogenesis (7, 8), mRNA splicing (3, 9–14), nucleocytoplasmic shuttling (15), stress granule formation (16, 17), and translation (18–20) in a variety of cell types. Most recently, TDP-43 has garnered attention for its involvement in several neurodegenerative diseases, particularly amyotrophic lateral sclerosis (ALS). In ALS, unknown neuronal stress triggers cause TDP-43 to form insoluble proteotoxic aggregates in the cytoplasm that ultimately drive motor neuron loss (21). The extent to which virus infection stress might influence TDP-43 function and whether TDP-43 potentially impacts virus infection biology beyond HIV is incompletely understood and understudied.

A limited number of molecular virus-host interactions involving TDP-43 have been reported, suggesting that TDP-43 may play a role during virus infection. Enterovirus infection was shown to cleave TDP-43 into shorter, more aggregate prone isoforms leading to the formation of TDP-43 inclusion bodies *in vivo* (22–24). Similarly, a viral protein in Theiler’s murine encephalomyelitis virus was shown to induce formation of cytoplasmic TDP-43 aggregates during infection (25). HSV-2 infection mildly upregulates TDP-43 in mouse spinal cords, but it is unclear if its intracellular localization is affected (26), and HSV-1 ICP0 reportedly promotes ubiquitination of TDP-43 (27). Finally, TDP-43 was found to bind and stabilize influenza A viral mRNAs, promoting infection (28). This raises the possibility that TDP-43 might impact the replication of other viruses, including neurotropic alphaherpesviruses like herpes simplex virus type-1 (HSV-1).

HSV-1 is a widespread human pathogen that establishes life-long latent infection in sensory ganglionic neurons (29). Reactivation of HSV-1 in response to neuronal stress leads to new infectious virus production and is generally associated with mild pathology such as cold sores, but can lead to life-threatening infection in the brain in neonates and immunocompromised individuals. While numerous host factors are effectively hijacked during HSV-1 infection to remodel the intracellular environment, neutralize host defenses, and facilitate virus productive replication, our understanding of how cell type specific roles for host factors, specifically in neuronal derived cells, might impact HSV-1 reproduction remains limited. In particular, how a pivotal neuronal stress responsive disease gene like TDP-43 that functions in RNA processing might influence HSV-1 gene expression and replication is unknown.

Here, we investigate the role of TDP-43 during HSV-1 infection of epithelial cells, fibroblasts and neurons. Following TDP-43 depletion in epithelial cells and primary fibroblasts, differences in productive HSV-1 replication were not detected. However, TDP-43 depletion was found to restrict virus replication in matured human HD10.6 cells, which are derived from dorsal root ganglia (DRG) sensory neurons. Depletion of TDP-43 in HD10.6 cells globally reduced virus gene expression and resulted in increased intron retention in mRNAs encoding ICP0, a critical master regulatory protein, and UL15, a key DNA packaging factor. Our data indicate that TDP-43 promotes global lytic gene expression and efficient splicing of key virus genes in a neuronal cell type-specific manner. It further suggests that normal TDP-43 levels are dispensable for viral replication in non-neuronal cells, but is required for efficient viral replication in a human DRG-derived neuronal cell line.

## RESULTS

### TDP-43 is dispensable for HSV-1 replication in epithelial cells and fibroblasts

To characterize the role of TDP-43 during HSV-1 infection, we first asked if the overall abundance of TDP-43 protein was changed upon HSV-1 infection. ARPE-19 retinal pigment epithelial cells and primary, normal human dermal fibroblasts (NHDF) were infected with HSV-1, and probed for TDP-43 protein levels by immunoblot at 9 hours post-infection (hpi). In both cell types, HSV-1 infection did not detectably alter TDP-43 abundance under these conditions (Fig 1A). To determine whether TDP-43 is required for efficient HSV-1 replication, ARPE-19s and NHDFs were transfected with two independent siRNAs targeting TDP-43 or a non-targeting control siRNA, and infected at low MOI with HSV-1 for 48 hpi. Whereas transfection with either TDP-43 siRNA resulted in a significant depletion of TDP-43 in ARPE-19s and NHDFs compared to non-targeting siRNA (Fig 1B), infectious virus titers were not detectably altered during this multi-cycle growth experiment (Fig 1C,D). Thus, TDP-43 protein abundance is not detectably changed by HSV-1 infection and TDP-43 depletion did not detectably interfere with productive HSV-1 replication in epithelial cells and primary fibroblasts.

**FIG 1.**
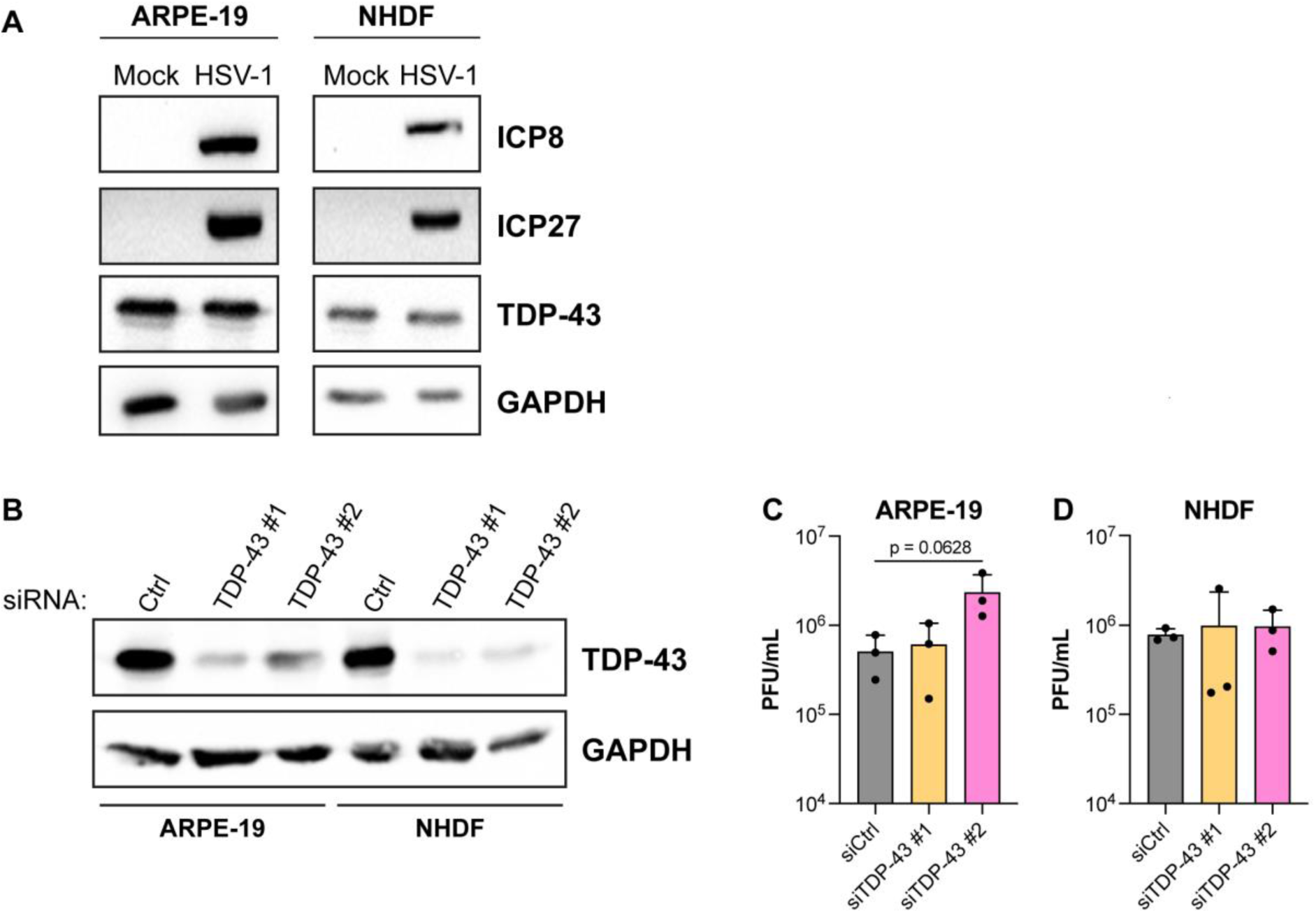
TDP-43 is dispensable for HSV-1 replication in epithelial cells and fibroblasts. (A) Immunoblots showing the abundance of TDP-43, ICP8 and ICP27 in uninfected (mock) or HSV-1 infected (MOI=3, 9 hpi) ARPE-19 cells or normal human dermal fibroblasts (NHDFs). GAPDH was used as a loading control. (B) Immunoblots confirming depletion of TDP-43 3 days following siRNA transfection using two independent siRNAs (TDP-43 #1, and TDP-43 #2) compared to a non-targeting control siRNA (Ctrl) in ARPE-19 cells and NHDFs. (C,D) Infectious virus titer at 48 hpi in the supernatant of ARPE-19 cells (C) and NHDFs (D), treated with a control siRNA or two siRNAs targeting TDP-43 for 72 h prior to infection with HSV-1 at MOI=0.0001.

### HD10.6 cells as a model for lytic HSV-1 infection of neurons

Although TDP-43 is a ubiquitously expressed RNA binding protein, TDP-43 associated pathology is predominantly limited to the central nervous system. Therefore, we investigated whether there is a unique role for TDP-43 during infection of neuronal derived cells compared to epithelial cells or fibroblasts. To model HSV-1 infection in neurons, we used matured HD10.6 cells. This cell line was derived from fetal human dorsal root ganglia (DRG) – a natural target cell for HSV-1 infection *in vivo* - and is immortalized through tetracycline regulated v-myc overexpression (30). Upon maturation using neuronal growth factors and doxycycline, HD10.6 cells express neuronal transcription factors, develop a neuronal morphology including connecting axons, and exhibit nociceptive properties. In addition, these cells can reportedly be rapidly matured into functional neurons that are permissive to HSV-1 infection (31). We confirmed that upon maturation, HD10.6 cells express elevated β3-tubulin and TrkA compared to proliferating controls, form dense neuronal networks, and can be infected with HSV-1 leading expression of viral lytic genes of all kinetic classes and ultimately the production of cell-associated infectious virus (Fig 2).

**FIG 2.**
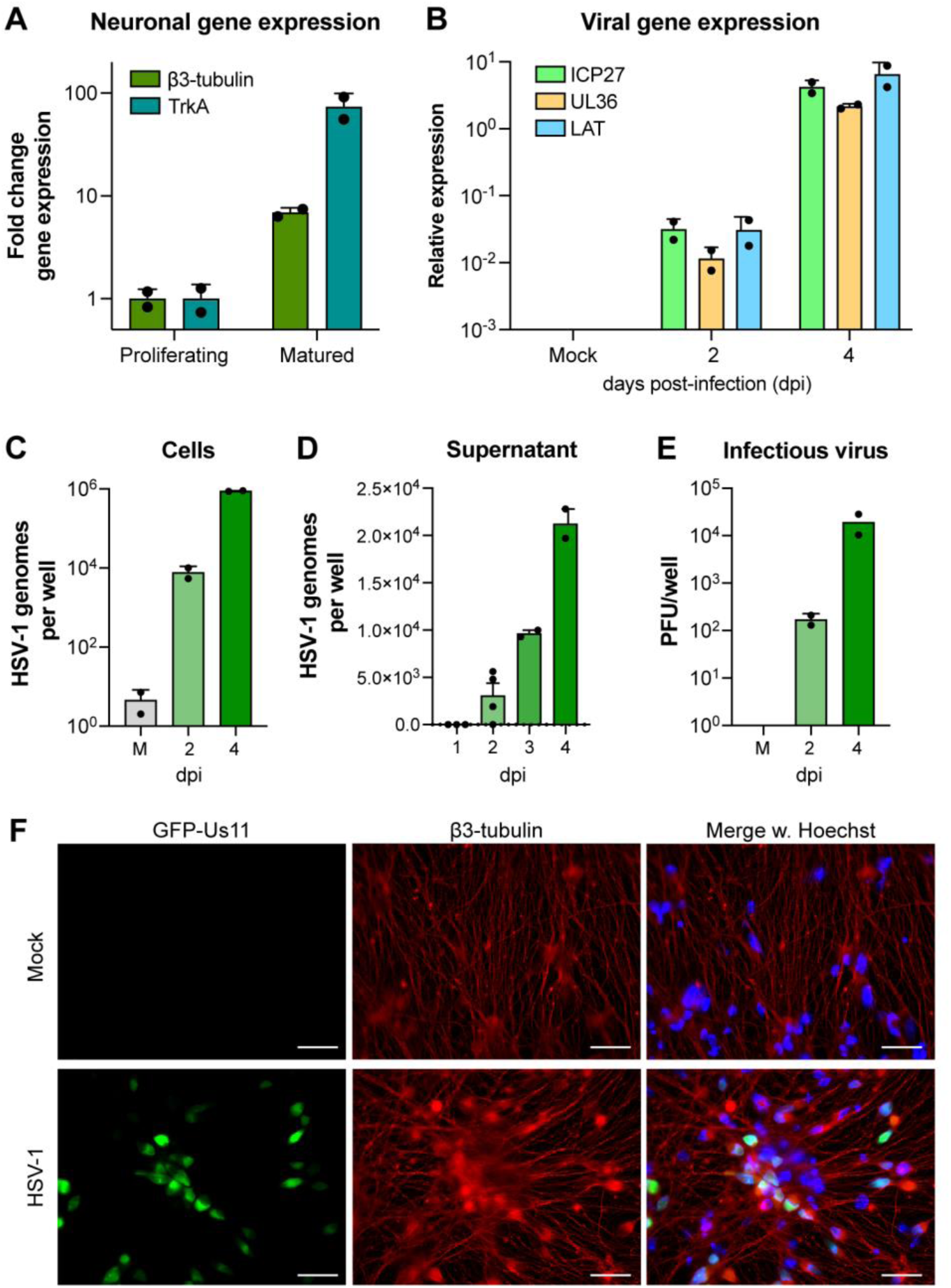
Validation of HD10.6 maturation and HSV-1 infection. HD10.6 cells were maturated for 10 days prior to infection with HSV-1 GFP-Us11 at MOI=0.1 for up to 4 days. Each data point represents an individually matured well. (A) Fold change of neuronal gene expression (β3-tubulin and TrkA) for proliferating or matured HD10.6. (B) Relative viral gene expression (ICP27, UL36, LAT) for HD10.6 cells uninfected (Mock), or infected for 2 or 4 days. (C,D) Absolute UL36 genome copies in HD10.6 cells (C) or their supernatant (D). (E) Infectious virus titer in HD10.6 cells. Titration of supernatant did not yield any detectable infectious virus. (F) Indirect immunofluorescence for HSV-1 GFP-Us11 (green) infected HD10.6 for 4 days, stained for β3-tubulin (red) and counterstained nuclei with Hoechst 33342 (blue). Epifluorescence imaging, magnification 63x and scale bar = 20 μM.

### TDP-43 depletion restricts HSV-1 replication in a multicycle growth curve in HD10.6 cells

We first asked if HSV-1 infection of HD10.6 cells alters TDP-43 mRNA and protein abundance. While a small but not statistically significant increase in TDP-43 mRNA was detected by RT-qPCR in HSV-1 infected compared to uninfected (Mock) HD10.6 cells (Fig 3A), changes in TDP-43 protein levels were not readily observed (Fig 3B) in agreement with findings in non-neuronal cells (Fig 1A). To determine if TDP-43 influences HSV-1 infection in HD10.6 cells, TDP-43 was depleted using transduction of two independent lentiviruses delivering shRNAs targeting TDP-43 or a non-targeting control shRNA (shNTC). Compared to cultures transduced with shNTC, TDP-43 protein abundance was substantially reduced by similar levels using either of the two TDP-43 specific shRNAs (Fig 3C, D). Since HD10.6 cells do not exhibit cytopathic effect upon HSV-1 infection, we used an HSV-1 reporter that expresses eGFP fused to the N terminus of the *true late* protein Us11 (HSV-1 GFP-Us11) allowing virus replication to be readily monitored in infected cultures (32). Notably, depletion of TDP-43 in HD10.6 cells resulted in a marked reduction in the accumulation of GFP expressing (GFP+) cells at 3 days post-infection (Fig 3E, F). Accordingly, significantly less – approximately 15-fold – infectious virus was produced as determined by plaque assay from HD10.6 cells in which TDP-43 was depleted compared to the non-targeting control lentivirus (Fig 3G). In accordance with reduced viral replication, we confirmed that expression of essential viral lytic genes representing *immediate-early* (ICP27) or *late* (UL36) kinetic classes was decreased in HD10.6 cells depleted for TDP-43 compared to cultures transduced with a non-targeting shRNA lentivirus (Fig 3H). Together, our data shows that while TDP-43 depletion does not detectably interfere with viral replication in epithelial cells and fibroblasts, it restricts HSV-1 gene expression and infectious virus production in a human DRG-derived cell line. This raises the possibility that TDP-43 is differentially required for HSV-1 replication in HD10.6 cells, but dispensable for productive virus growth in non-neuronal cells.

**FIG 3.**
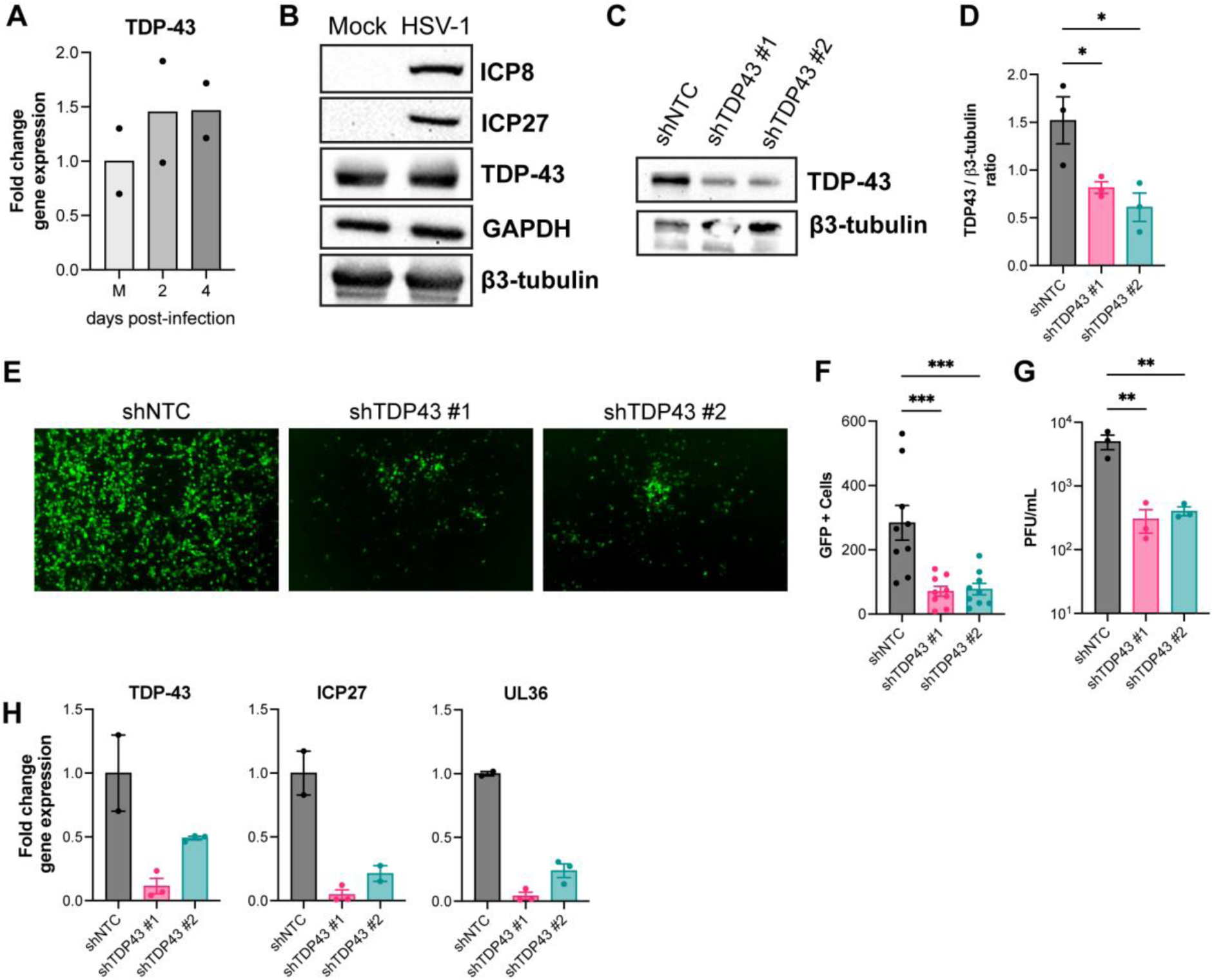
TDP-43 knockdown restricts HSV-1 replication in a multicycle infection of HD10.6 cells. (A) Fold change TDP-43 gene expression in uninfected (M) or HD10.6 cells infected for 2 or 4 days at MOI=0.1 with HSV-1 GFP-Us11 virus. (B) Immunoblots showing the abundance of TDP-43, ICP8 and ICP27 in uninfected (Mock) or HSV-1 infected (MOI=30, 24 hours) HD10.6 cells. GAPDH and β3-tubulin were used as loading control. (C-H) HD10.6 cells were transduced with lentiviruses expressing non-targeting shRNA (shNTC) or two independent TDP-43 targeting shRNAs (#1 or #2) for 3 days and subsequently infected with HSV-1 GFP-Us11 at MOI=0.5 (C-G), or MOI=0.1 (H) for 3 days, n=2-3 individual wells per condition. (C) Representative immunoblots for TDP-43 abundance with β3-tubulin as loading control with (D) intensity quantification of immunoblots, p < 0.05 by one-way ANOVA. (E) Representative fluorescent images of HSV-1 GFP-Us11 infected shRNA-transduced HD10.6 at 3 days post-infection, with (F) quantification of GFP+ cells (n=3 images per well), p < 0.001 by one-way ANOVA. (G) Cell-associated virus titer as determined by plaque assay, p < 0.01 by one-way ANOVA. (H) Fold change in TDP-43 and viral (ICP27, UL36) gene expression in HSV-1 GFP-Us11 infected shRNA-transduced HD10.6 at 3 days post-infection.

### HSV-1 viral lytic gene expression is limited upon TDP-43 knockdown

HSV-1 lytic replication is broadly divided into three kinetic phases: *immediate-early, early* and *late*. To establish when in the viral life cycle TDP-43 depletion impacts HSV-1 gene expression, we performed a single-cycle infection. In our hands, HD10.6 cells were relatively resistant to HSV-1 infection, and only at an MOI of at least 30 – based upon virus titered on permissive cells – the majority of the cells were infected at 24 hpi, and viral titers appeared to plateau (Fig 4A, B). Notably, even under high MOI conditions and in a restricted time frame, TDP-43 depletion significantly reduced the number of GFP+ cells (Fig 4C, D) and reduced infectious virus production by approximately four-fold (Fig 4E), suggesting TDP-43 depletion restricts HSV-1 replication within a single replication cycle. Evaluating expression of representative virus genes from different kinetic classes by RT-qPCR revealed that viral lytic gene expression was broadly repressed by TDP-43 knockdown (Fig 4F). This included significant reductions in the abundance of mRNAs encoding the critical *immediate-early* (IE) regulatory proteins ICP0 and ICP4. Indeed, TDP-43 depletion effectively reduced representative HSV-1 IE mRNAs (ICP0, ICP4, ICP27) abundance in infected cells treated with cycloheximide, which inhibits *de novo* protein synthesis and only allows virus *immediate-early* gene transcription (Fig 4G). Together, this suggests that TDP-43 might control HSV-1 infection through changes to both virus and the host.

**FIG 4.**
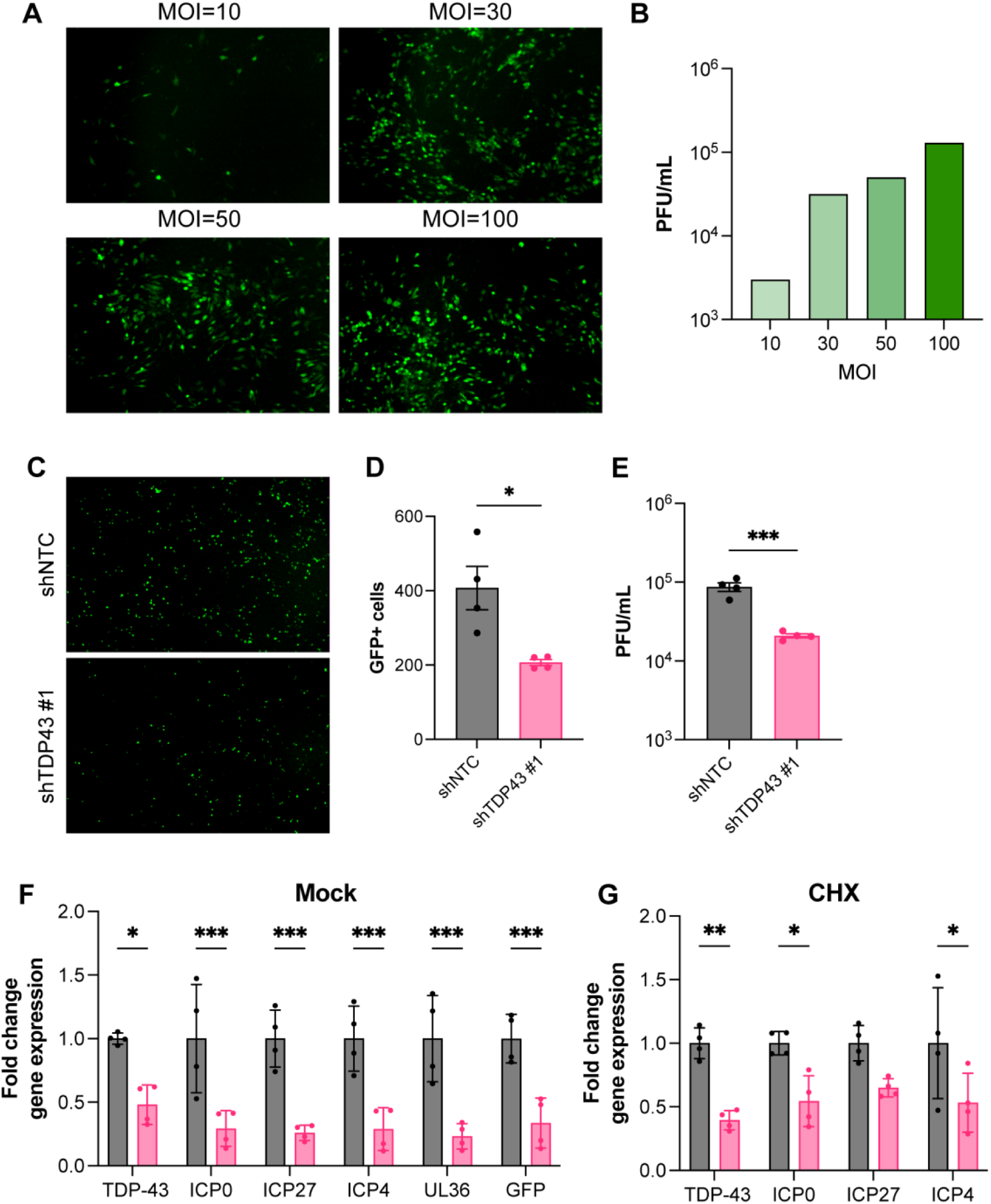
TDP-43 depletion limits HSV-1 gene expression during high MOI infection of HD10.6 cells. (A) Representative images of matured HD10.6 infected with HSV-1 GFP-Us11 with MOIs 10, 30, 50 or 100 at 24 hpi and (B) Infectious virus titer retrieved from HD10.6 cells at indicated MOIs. (C-G) HD10.6 cells were transduced with lentiviruses expressing non-targeting shRNA (shNTC) or TDP-43 targeting shRNA #1 for 3 days and subsequently infected with HSV-1 GFP-Us11 at MOI=30 for 24 hours, n=4 individual wells per condition. (C) Representative fluorescent images of HSV-1 GFP-Us11 infected shRNA-transduced HD10.6 cells at 24 h post-infection, with (D) quantification of GFP+ cells, p < 0.05 by Student’s t-test. (E) Cell-associated virus titer as determined by plaque assay, p < 0.001 by Student’s t-test. (F-G) Fold change in TDP-43 and viral (ICP0, ICP27, ICP4, UL36 and eGFP) gene expression in HSV-1 GFP-Us11 infected shRNA-transduced HD10.6 cells at 24 days post-infection, treated with DMSO (F) or cycloheximide (G), n=4 per condition. * p < 0.05, ** p < 0.01, or *** p < 0.001 by two-way ANOVA.

### TDP-43 depletion results in a genome-wide reduction of viral gene expression

In uninfected cells, several studies have shown a myriad of effects on TDP-43 depletion on RNA biology, including direct changes to the host transcriptome through differential splicing and poly-adenylation. To more globally assess whether specific HSV-1 RNAs are changed upon TDP-43 depletion, we employed an unbiased transcriptome-level sequencing approach. Nanopore direct RNA-sequencing (dRNA-seq) is a powerful technique that allows for sequencing of full length RNAs, capturing transcription start (TSS) and termination sites (TTS) as well as splice junctions (33, 34). This is particularly useful when applied to highly dense viral genomes harboring multiple polycistronic transcription units that can be difficult to deconvolute using short-read sequencing approaches (34–36). In accordance with restricted viral replication, shRNA-mediated depletion of TDP-43 in HD10.6 cells resulted in the detection of proportionally fewer (4-fold) viral reads by dRNA-seq (Fig 5A, B). Next, we quantified the relative abundance of each viral transcript in both conditions (transcripts per million, TPM), and calculated the ratio of knockdown over control. The TPM ratio was below 1 for all viral genes, with no significant difference between genes considered essential or non-essential for viral replication (Fig 5C). Furthermore, a significant difference in depletion between viral transcripts of different kinetic classes was not observed (Fig 5D). Together, this suggests that TDP-43 depletion leads to a genome-wide restriction of viral lytic gene expression.

**FIG 5.**
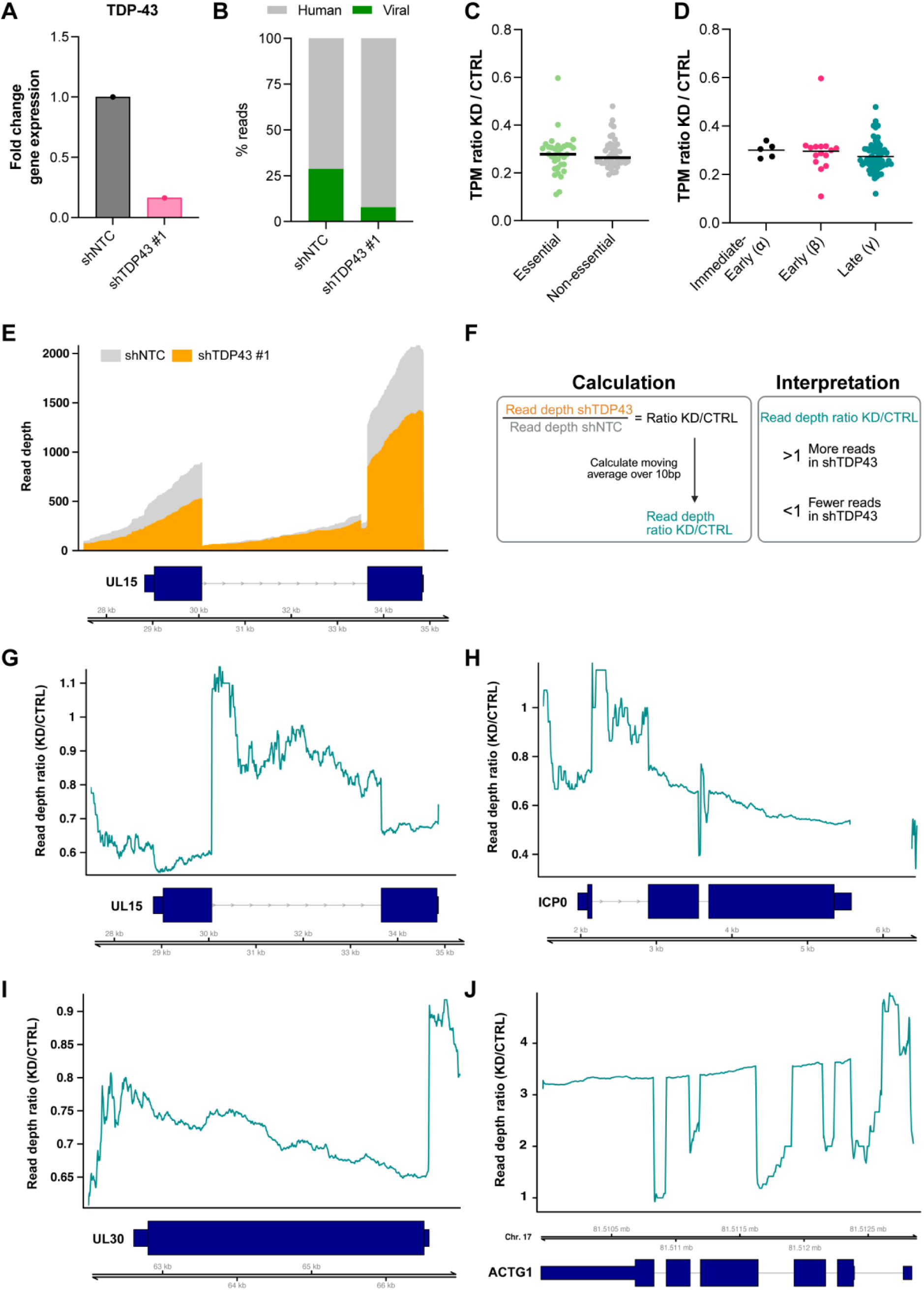
TDP-43 loss leads to global repression of HSV-1 and increased intron retention in HD10.6 cells. HD10.6 cells were transduced with lentiviruses expressing non-targeting shRNA (shNTC) or TDP-43 targeting shRNAs #1 for 3 days and subsequently infected with HSV-1 GFP-Us11 at MOI=30 for 24 hours. (A) Fold change in TDP-43 gene expression. (B) Percentage of viral and human reads in nanopore dRNA-seq. (C,D) Ratio of transcript per million (TPM) of shTDP43 #1 (KD) over shNTC (CTRL) for each individual viral transcript, with (C) categorized for essential and non-essential viral genes and (D) kinetic classes. (E) Strand-specific raw read depth (y-axis) over the UL15 locus in shNTC (grey) and shTDP43 #1 (orange) samples. (F) Schematic of calculation of the moving average (window size=10b) of the coverage ratio of KD over CTRL plotted as teal lines in (G-J) and the interpretation. (G-J) Moving average of read depth ratio of KD over CTRL plotted as a teal line, with (G) depicting UL15, (H) ICP0 and (I) UL30 and human gene ACTG1 (J). (E,G-J) Dark blue boxes indicate RNA structures, with thin boxes representing untranslated regions (UTR), wide boxes coding sequence (CDS) and arrowed lines introns. Double black lines represent the HSV-1 genome (E, G-I) or human genome (J).

### Evidence for enhanced intron retention in TDP-43 knockdown cells

To determine whether TDP-43 influences the architecture of the HSV-1 transcriptome through alternative TSS or TTS usage, we extracted the start and end coordinates for all viral reads and filtered out all with an abundance of less than 10 reads, to yield frequency estimates for the 5’ and 3’ ends of polyadenylated viral transcripts. In doing so, we did not observe any unique or differentially used putative TSS or TTS (pTSS or pTTS) for HSV-1 transcripts in TDP-43 depleted HD10.6 cells, suggesting that TDP-43 does not detectably influence transcription initiation or termination during HSV-1 infection. Additionally, in human neurons, TDP-43 has been shown to control splicing of specific host genes through intron retention or cryptic exon inclusion, thereby altering transcript stability (3, 9, 10, 13, 14). Whereas the majority of mammalian genes are spliced, only five HSV-1 transcripts consistently undergo splicing, namely: ICP0, UL15, Us1, Us12 and LAT. Interestingly, when we plotted the read depth across UL15 for both control and TDP-43 depleted HD10.6 cells we observed a marked reduction of coverage in TDP-43 depleted cells in the exons, but similar abundance in the intronic region (Fig 5E). To visualize this better, we calculated the read-depth ratio and plotted the moving average over 10 bases. Above 1, this ratio indicates a higher coverage in the TDP-43 depleted sample than the control at that genomic location, whereas below 1 the coverage is higher for the control sample (Fig 5F). Interestingly, the coverage ratio ranged from 0.6-0.7 in both UL15 exons but was increased sharply across the intron ranging from 0.8-1.1 (Fig 5G). This suggests that upon TDP-43 depletion, UL15 RNAs are less efficiently spliced. We found similar evidence of intron retention in the first ICP0 intron (Fig 5H), but not for the Us1/Us12 intron implying a level of selectivity. An unspliced control gene UL30 did not exhibit similar variation in read depth ratio throughout the gene body, suggesting this is a unique signature of spliced genes (Fig 5I). As the primary LAT transcript is very unstable and the intron is not polyadenylated, we were unable to retrieve any reads corresponding to LAT. Together, this data demonstrates that in addition to a global control of viral gene expression, TDP-43 also promotes more efficient splicing of two viral genes, one of which encodes the global IE regulatory protein ICP0.

### Intron retention in UL15 is dependent on the relative abundance of TDP-43

To confirm enhanced intron retention in ICP0 and UL15 mRNAs, we designed primer sets that can distinguish between spliced and intronic variants of these mRNAs (Fig 6A). We observed a slight increase (14%) in the relative abundance of the first ICP0 intron upon TDP-43 knockdown (Fig 6B). However, under CHX treatment, which inhibits protein synthesis and selectively allows only IE gene transcription, the abundance of ICP0 intron-containing transcripts increased by 2.5-fold HD10.6 cells depleted for TDP-43 compared to control. (Fig 6C). This could reflect that many intronic transcripts are degraded by nonsense-mediated decay (NMD) machinery in the cytoplasm, which is a translation dependent mechanism. Alternatively, this could indicate a viral factor – potentially ICP27 which can limit mRNA splicing (37–40) – reducing the abundance of intron containing transcripts. Similarly, we observed a slight increase in intronic UL15 RNAs upon TDP-43 depletion (Fig 6D), that significantly correlated to the extent of TDP-43 knockdown in the same sample (Fig 6E). Together this data shows that UL15 splicing efficiency in HD10.6 cells is influenced by TDP-43 abundance.

**FIG 6.**
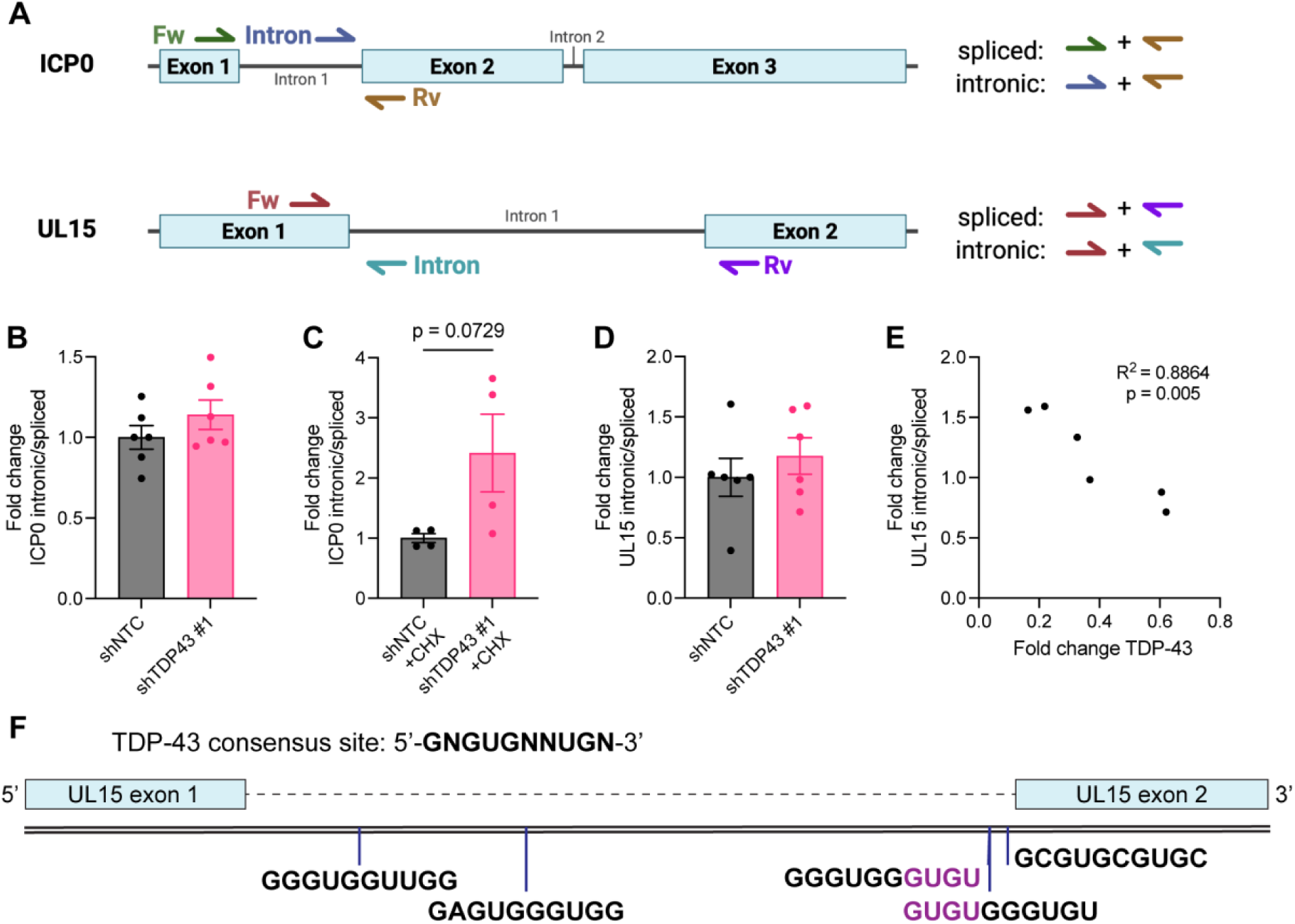
TDP-43 loss in HD10.6 cells mildly increases detection of intron containing ICP0 and UL15 mRNAs by RT-qPCR. (A) Schematic representation of ICP0 and UL15 mRNA regions, indicating approximate locations of forward, reverse and intronic primers. For ICP0, forward primers distinguish between spliced and intronic mRNAs, whereas reverse primers distinguish between spliced and intronic UL15. (B-E) HD10.6 cells were transduced with lentiviruses expressing non-targeting shRNA (shNTC) or TDP-43 targeting shRNAs #1 for 3 days and subsequently infected with HSV-1 GFP-Us11 at MOI=30 for 24 hours. (B,C) Fold change of the ratio of intronic over spliced ICP0 mRNAs in untreated (B) or CHX-treated (C) HSV-1 infected HD10.6 cells. (D) Fold change of the ratio of intronic over spliced UL15 mRNAs. (E) Pearson correlation between UL15 intronic ratio and the relative abundance of TDP-43 mRNAs. (F) Schematic representation of UL15 genomic region, indicating locations of consensus TDP-43 binding sites (blue lines) in the UL15 intron. mRNA sequences for each consensus binding site (5’->3’) are given in black, and purple letters indicate shared nucleotides between sites 3 and 4.

## DISCUSSION

A hallmark of several prominent neurodegenerative disorders is proteotoxic accumulation of key neuronal proteins, such as amyloid-β, tau, α-synuclein, TDP-43 (41–43). Recent studies implicate herpesvirus reactivation in the onset of these neurodegenerative disorders (44–47), though little is known about virus-host interactions in neuronal cell types. Here, we have shown differential requirement for TDP-43 expression during lytic HSV-1 infection in the human neuronal derived HD10.6 cell line compared to epithelial cells and primary fibroblasts. We show that depletion of TDP-43 restricts viral replication in HD10.6 derived neurons during both single-cycle and multi-cycle HSV-1 infection, through generalized transcriptional repression and enhanced intron retention in key virus-encoded mRNAs. As such, this study demonstrates the importance of using neuronal derived cell models to uncover cell type specific regulatory mechanisms.

TDP-43 is an RNA-binding protein that is expressed by all cell types, but of these, neurons are considered uniquely sensitive to TDP-43 depletion, mutations or mislocalization. We showed that TDP-43 depletion specifically impairs HSV-1 replication in a model of human DRG-derived neurons (Figure 3,4), but not in epithelial cells or fibroblasts (Figure 1). Limited research into neuron specific responses and requirements to HSV-1 infection suggests that neurons have a reduced antiviral response, and are uniquely sensitive to depletion of specific proteins (48–51). The requirement of a neurotropic virus, exemplified here by HSV-1, for a host TDP-43 protein involved in neuronal homeostasis, highlights how viruses repurpose host processes for their own benefit.

Despite the fact that TDP-43 was first identified as restricting HIV-1 infection, few studies have addressed its role in promoting or antagonizing replication of other viruses. In this study, we show that HSV-1 mRNA accumulation is reduced upon TDP-43 depletion in HD10.6 cells, even in the absence of viral protein synthesis. This raises the question whether TDP-43 could directly regulate viral gene expression. In HIV-1 infected cells, TDP-43 binds to the TAR DNA sequence and limits its transcription (4), and stabilizes HDAC6 expression – a key HIV restriction factor - further inhibiting viral replication (52, 53). More recently, TDP-43 was shown to be recruited by the viral polymerase of influenza A virus, bind and stabilize viral mRNAs (28). Notably, TDP-43 depletion modestly impairs influenza viral growth, similar to our observations for HSV-1 in HD10.6 cells. It remains to be explored if HSV-1 mRNAs in HD10.6 cells are stabilized through direct TDP-43 binding, or if TDP-43 controls HSV-1 transcription through interaction with RNA Pol II or viral DNA. Notably, about half of the viral mRNAs contain at least one TDP-43 minimal consensus sequence (54), suggesting that TDP-43 interactions with viral mRNAs are possible. TDP-43 reportedly autoregulates its abundance through a negative feedback loop by binding to the 3’UTR of TDP-43 mRNAs (55, 56), and similarly regulates the stability of other mammalian mRNAs through binding to the 3’UTR (11, 57, 58).

A well-characterized function of TDP-43 in neurons is ensuring correct splicing, through repression of cryptic exons and retained introns in neuronal genes (3, 9–11, 19, 59). Using dRNA-seq to interrogate complete transcript structures in HSV-1 infected cells, we showed that TDP-43 depletion leads to a modest enhancement of intron retention in two viral genes, RL2 (ICP0) and UL15 (TRM3), that are important for efficient viral replication (60). Notably, intron containing viral mRNAs – specifically those of ICP0 and UL15 – are thought to accumulate in the nucleus as they are less efficiently exported to the cytoplasm by the viral protein ICP27 (38, 61, 62). Enhanced intron retention resulting from TDP-43 depletion could thus alter the availability of spliced mRNAs for translation and impair efficient replication. Specifically, an early deficit in ICP0 abundance could in part contribute to the global reduction in gene expression upon TDP-43 depletion, as antiviral defenses might not be effectively countered. The increased abundance of intron-containing ICP0 transcripts in the absence of viral protein synthesis (Fig 6C) suggests the mechanism by which TDP-43 regulates viral mRNA splicing is distinct from the ICP27-dependent mechanisms of other host splicing factors (37, 40, 63, 64). Interestingly, a manual search revealed that UL15 pre-mRNA contains 5 minimal TDP-43 binding sites (54) throughout the intron (Fig 6F), raising the possibility that UL15 splicing might be regulated similarly to host genes where TDP-43 represses introns through direct binding (11).

In summary, we showed that the critical neuronal disease gene TDP-43 specifically regulates replication of a common, widespread neurotropic HSV-1 in neuron-derived cells but not epithelial cells or primary fibroblasts. Our work highlights the importance of investigating virus-host interactions in multiple relevant model systems, as proteins can have cell-type specific functions. Furthermore, as human alphaherpesvirus reactivation is increasingly linked to the onset of Alzheimers’ disease, this study raises the question whether HSV-1 could also be a contributing factor in other neurodegenerative disorders such as ALS. Future studies should leverage neuronal culture systems to determine whether HSV-1 induced infection stress or virus functions influence neurodegeneration resulting from TDP-43 depletion or aggregation.

## MATERIALS AND METHODS

### Cell culture

Normal human dermal fibroblasts (NHDFs) were cultured in DMEM supplemented with 5% heat-inactivated FBS (HI-FBS, Gibco) and 1% penicillin-streptomycin (PS, Lonza). Human retinal pigmented epithelial ARPE-19 cells were cultured in DMEM:F12 (Gibco) supplemented with 10% HI-FBS and 1% PS, 15 mM HEPES and 1 mM sodium pyruvate (Gibco). 293T Lenti-X cells (Takara Bio) were maintained in DMEM supplemented with 10% HI-FBS, 1% PS. Vero cells were maintained in DMEM supplemented with 5% calf-serum and 1% PS.

### Culture and Maturation of HD10.6 cells

Proliferating HD10.6 cells (a gift of Anna Cliffe, University of Virginia School of Medicine) were maintained on 17 μg/mL fibronectin in PBS pre-coated T-75 flasks in Proliferation Media: Advanced DMEM/F12 (Gibco), supplemented with 1x Neurocult SM1 (STEMCELL Technologies), 10 ng/mL prostaglandin E1 (Sigma), 2mM L-glutamine (Gibco), 1x Primocin (Invivogen) and fresh 0.5 ng/mL bFGF (PeproTech). For maturation, HD10.6 were plated at 25.000 cells/cm2 in Proliferation Media on plates precoated with 50 μg/mL poly-L-ornithine hydrobromide (Sigma) in 0.5 M borate buffer pH 8.5 (Boston Bioproducts), and 1 μg/mL fibronectin. One day after plating, media was replaced for Maturation +/+/+ Media: Neurobasal (Gibco), supplemented with 1x Neurocult SM1, 2 mM L-glutamine, 1x Primocin, 1 μg/mL doxycycline, 50 ng/mL NGF (Alomone Labs), 25 ng/mL CNTF, GDNF and NT-3 (all Peprotech).

### HSV-1 stocks and infections

Working stocks of wild type HSV-1 strain KOS and HSV-1 GFP-Us11 strain Patton were propagated on Vero or ARPE-19 cells respectively. Virus stocks, supernatants (NHDF, ARPE-19) or cell-associated virus (HD10.6) were titered using plaque assays on Vero cells. For experiments, NHDF, ARPE-19 and Vero cells were infected for 1 hour in low serum (1%) media, or for HD10.6 cells for 2 hours in Maturation Base Media (Neurobasal with L-glutamine).

### Transfection of siRNAs

NHDFs and ARPE-19s were seed at approximately 30% confluency, and transfected with 20 nM siRNA using Lipofectamine RNAiMax (Life technologies) in Optimem (Gibco) 3 days prior to infection. siRNAs used in this study were purchased from Sigma: siTDP-43-#1 (SASI-Hs01-0037054) and siTDP-43-#2 (SASI-Hs01-0037055). AllStars Negative Control siRNA (Qiagen) was used as the non-targeting siRNA control.

### Lentiviral generation and transduction

Lentiviruses were generated by transfecting equimolar ratios of monomeric pMD2.G (VSV-G, derived from Addgene #12259), psPAX2 (Addgene #12260) and transfer plasmids into subconfluent 293T Lenti-X cells (Takara Bio) using a 3:1 ratio of linear polyethylenimine MW 25000 (Polysciences). Media was replaced 1 day later for Optimem (Gibco), and supernatant was collected 3 days post-transfection and passed through 0.45 μm PVDF filter before aliquoting and storage at -80°C. Lentiviral transfer plasmids used in this study were: pLKO.1-puro Non-Target shRNA control plasmid as negative control, TRCN0000016040 (shTDP43-#1) and TRCN0000016041 (shTDP43-#2), all from Sigma. To transduce HD10.6 cells, approx. 30 μL lentiviral stock per well was used (24-wells format), and incubated in half the original culturing volume of Maturation +/+/+ Media overnight without polybrene. The next day, lentivirus containing media was removed and replaced with fresh Maturation +++ media. Cells were used for assays 3-4 days post-transduction.

### Immunofluorescence

HD10.6 cells were plated for maturation on glass coverslips, and infected after 10 days of maturation. Cells were then fixed with 4% paraformaldehyde, permeabilized for 10 min with 0.1% Triton X-100 in PBS, and blocked with 5% normal goat serum diluted in PBS. Coverslips were incubated with rabbit anti-β3-tubulin (1:2000, 802001, BioLegend) primary antibody in PBS overnight. The next day coverslips were washed and incubated for 1hr at room temperature with AF555-conjugated goat anti-rabbit secondary antibody in PBS (1:500, Thermo Fisher Scientific). Finally, the coverslips were washed and incubated with Hoechst 33342 in PBS (20 μM, Life Technologies) for 5 minutes, washed and mounted using Fluorshield mounting media (Sigma). Stained coverslips were analyzed using a Leica DM5000B epifluorescence microscope with 63x objective lens with oil. Images were adjusted for brightness and contrast using Photoshop CC 2025 software (Adobe).

### Immunoblots

Cells were lysed using 1x Laemmli buffer and samples were incubated at 95°C for 10 min. Proteins were separated on a 10% bis-acrylamide gel, and transferred onto nitrocellulose membranes. Membranes were blocked using 5% non-fat dry milk in TBS-T for 1hr, and incubated with primary antibodies diluted in TBS-T + 3% BSA + 0.02% NaN3 overnight at 4°C with agitation. Primary antibodies used were rabbit anti-TDP-43 (10782-2-AP, Proteintech), rabbit anti-GAPDH (2118, Cell Signaling Technologies), rabbit anti-β3-tubulin (802001, BioLegend), mouse anti-ICP8 (ab20193, Abcam), mouse anti-ICP27 (ab31631, Abcam). The next day, membranes were washed thrice with TBS-T and incubated with 1:5000 of HRP conjugated secondary antibody (Cytiva) for 1 hour at RT in TBS-T + 5% milk. Membranes were then washed thrice with TBS-T and chemiluminescent signal was visualized using Pierce ECL Western Blotting kits at Regular or Pico strengths (ThermoFisher Scientific) on an iBright CL1000 imaging system.

### RNA isolation and cDNA synthesis

Cells were harvested in Trizol (Thermo Fisher Scientific), mixed with 1:5 chloroform and centrifuged for 15 min at 12,000xg at 4°C. From the aqueous phase, RNA was isolated using the RNeasy Mini kit (Qiagen) according to manufacturer’s instructions. Residual DNA was removed using the Turbo DNA-free kit (Ambion). To generate cDNA, Turbo-DNAse-treated RNA was reversed transcribed with Superscript IV reverse transcriptase (RT+) and random hexamers (ThermoFisher Scientific). The same cDNA synthesis reaction in the absence of reverse transcriptase (RT-) was performed to generate negative controls.

### Quantitative PCR analysis

Quantitative PCR (qPCR) was performed in duplicate on RT+ cDNA using 0.5 μm of each primer (IDT), 1x SsoAdvanced Universal SYBR Green Supermix (1725275, Biorad) on a Bio-Rad IQ5 thermal cycler, and RT-cDNA was run for select targets to exclude residual DNA carryover. An annealing temperature of 58°C was used for all targets. Melt curves were inspected for the presence of secondary peaks prior to data analysis. Primer sequences used in this study are given in Table 1 below. Relative expression was defined as 2−(Cq-value target gene − Cq-value housekeeping) and fold change as by normalizing relative expression to the average of control samples. Absolute HSV-1 genome copies were determined by using a 10-fold serial dilution of the UL36 amplicon (gBlock, IDT) in 20 ng/μL salmon sperm DNA (Invitrogen).

**Table 1.**
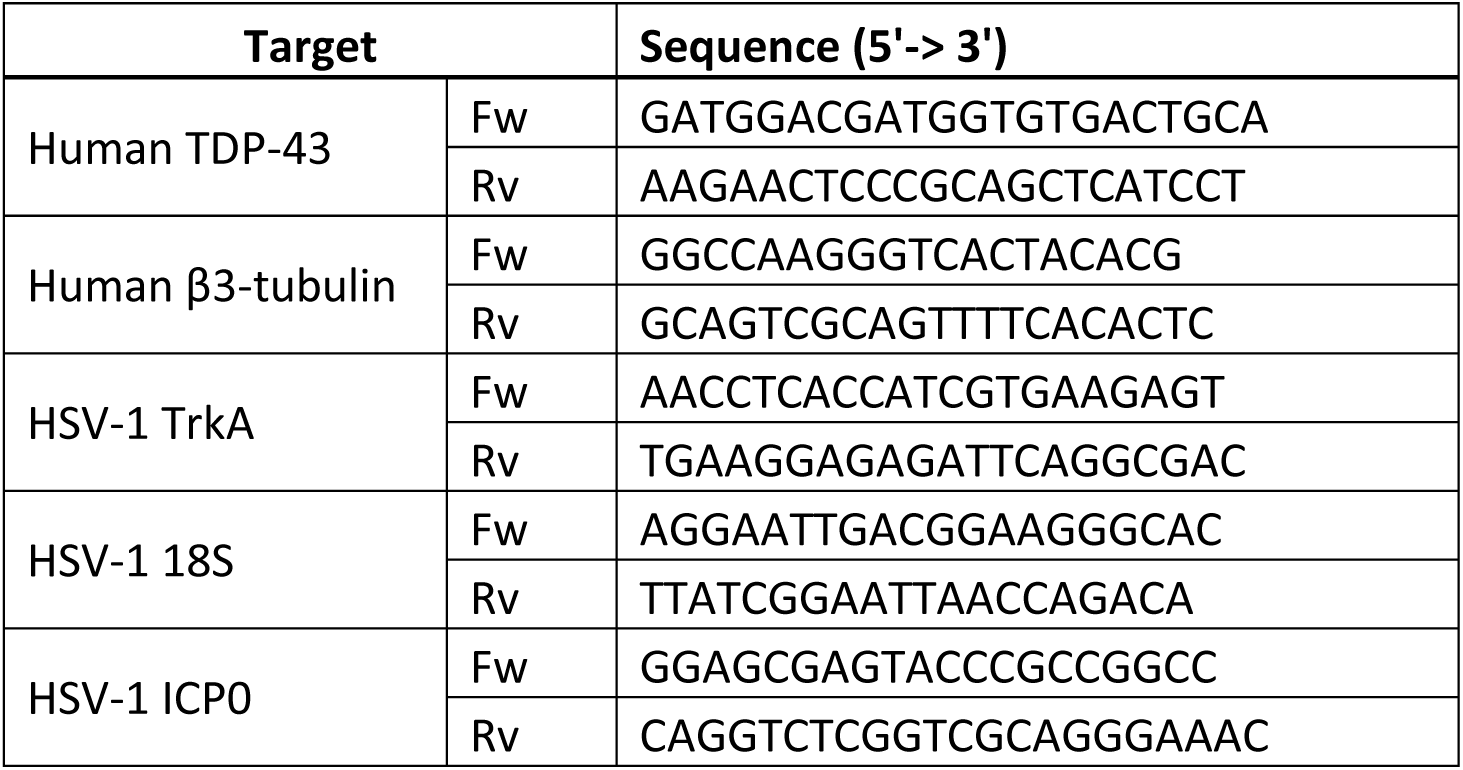
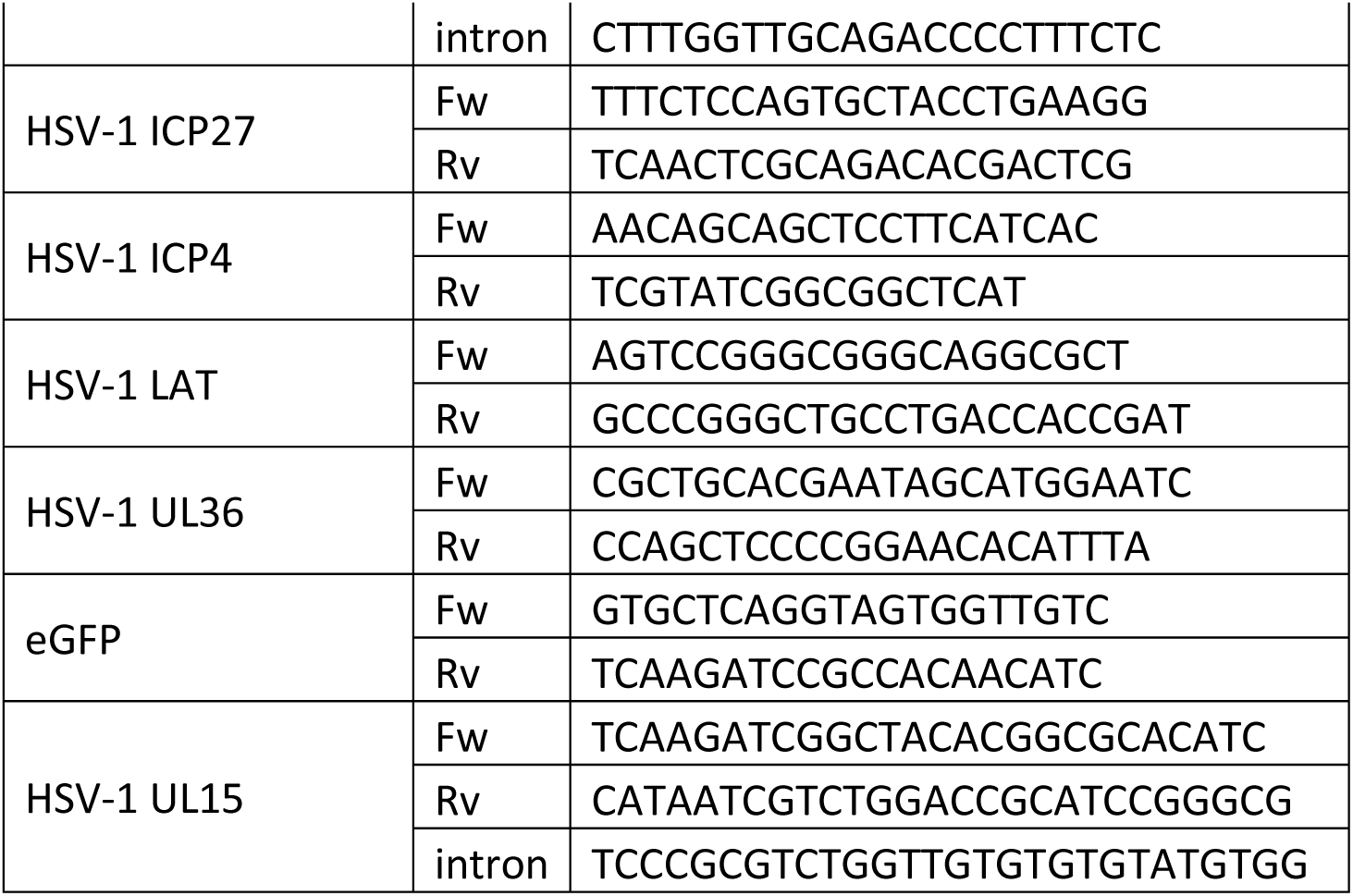
Primers used in this study.

### Nanopore dRNA-sequencing

To generate dRNA-sequencing libraries, poly(A) RNA was isolated using a Dynabeads mRNA purification kit (Invitrogen). 500 ng poly(A) served as input for library preparation using the direct RNA Sequencing (DRS) SQK-RNA004 kit. This resulted in 150 ng of library that was loaded onto FLO-MIN004RA flow cells. Sequencing was performed on a MinION Mk1B for 24 hours, yielding 1-2 million reads per sample.

### Nanopore bioinformatic analysis

Basecalling was performed with dorado v.0.8.1 (Oxford Nanopore Technologies), using a high accuracy model (rna004_130bps_hac@v5.0.0) with adapter trimming. FASTQ files were extracted from the unaligned BAM files using BEDtools v.2.30.0 (65). Reads were aligned to the human (hg38) or HSV-1 GFP-Us11 (MF959544.1) genomes using minimap2 v.2.24 (66) with spliced alignment settings (-ax splice) and a k-mer size of 14 (-k14). The resulting SAM files were filtered to exclude unmapped, secondary, and supplementary alignments (-F2308) using SAMtools v.1.16 (67), then converted to BAM format, sorted, and indexed.

For transcriptome alignment, FASTQ files were aligned to the HSV-1 GFP-Us11 and human transcriptomes (Gencode v47) using minimap2, with the setting -ax map -ont for Oxford Nanopore data and a minimum secondary-to-primary score ratio of 0.99 (-p 0.99). The resulting alignments were stored in SAM format. SAM files were filtered to retain primary alignments by removing supplementary, secondary and unmapped reads (-F2324). Subsequently, reads with hard clipping in the CIGAR string were removed. Only reads that start within 50 bp downstream of annotated start sites were retained. The filtered reads were converted into BAM files, sorted, and indexed with samtools.

Transcript abundance was quantified by extracting reference names from the sorted BAM file using samtools view, isolating the third column (reference name) with cut -f3, sorting entries alphabetically with sort, counting unique occurrences of each identifier using uniq -c, and exporting the results to a text file for downstream analysis.

To normalize differences in read depth, each transcript count was divided by the total number of mapped and filtered reads and multiplied by 1x106, denoted by transcripts per million (TPM). TPM values from TDP43-KD samples were divided by control samples to calculate a TPM KD/control ratio. Only locations with a minimum raw read depth of 10 were considered for downstream analysis.

pTSS and pTTS were determined through extracting the 5’ and 3’ end of each read using Rsamtools and locations with less than 10 counts were filtered out. pTSS and pTTS between samples were compared by plotting the abundance per location in each area of the genome. Coverage plots were generated using Genomic Ranges in RStudio, and gene models were generated from updated GFF3 files reflecting the HSV-1 GFP-Us11 transcriptome. To calculate coverage ratio the read depth for each base in shTDP-43 dataset was divided by the read depth in the shNTC dataset, and locations with a read depth below 10 in either dataset were excluded. To calculate the moving average and smoothen out local variations in alignment, the stats::filter function in R was used, with a window size of 10 bases.

### Statistical analysis

Figures show individual data points or mean +/- SEM, all statistical analyses were performed with Graphpad Prism 10 software using the statistical test indicated in the figure legends, with * = p<0.05, ** = p<0.01, *** = p<0.001. If no statistics are given, the difference was not significant.

## Data availability

All sequencing datasets generated as part of this study are available at the European Nucleotide Archive under the accession number PRJEB89368. Genome and transcriptome annotation files for HSV-1 GFP-Us11 are available on GitHub https://github.com/DeniseOhnezeit/HSV1-GFP-US11_Patton-reference-files.

## ACKNOWLEDGEMENTS

The authors would like to thank Anna Cliffe (University of Virginia School of Medicine) for sharing HD10.6 cells and protocols. S.E.B was supported by a Rubicon Postdoctoral Fellowship from the Nederlandse Organisatie voor Wetenschappelijk Onderzoek (NWO ZonMw, Dutch Research Council) project 452022210. D.O. was supported by the Walter Benjamin Programme of the Deutsche Forschungsgemeinschaft (DFG, German Research Foundation), project 554758329. The work was also supported by the following grants from the National Institutes of Health R01-AI176335 and R01-AI170583.

